# NucMerge: Genome assembly quality improvement assisted by alternative assemblies and paired-end Illumina reads

**DOI:** 10.1101/483701

**Authors:** Ksenia Khelik, Alexander Johan Nederbragt, Geir Kjetil Sandve, Torbjørn Rognes

## Abstract

**Background:** In spite of the major breakthroughs in the second-generation sequencing technologies and the developments of a plethora of assemblers over the last ten years, the resulting genome assemblies may still be fragmented and contain errors. It is typical in genome projects with second-generation reads involved to run multiple assemblers with different parameters and choose the best assembly. However, such an approach is always a trade-off between the strengths and weaknesses of the assemblies. To exploit the advantages of different assemblers, an alternative approach that combines the best parts of several assemblies into one may be applied. The existing tools based on such an approach assist in elongation of assembly fragments and/or improvement of assembly accuracy. Though there has been progress with such a strategy, there is still room for improvement of the existing tools.

**Results:** We present NucMerge, a tool for improving genome assembly accuracy by incorporating information derived from an alternative assembly and paired-end Illumina reads from the same genome. The tool corrects insertion, deletion, substitution, and inversion errors and locates different inter- and intra-chromosomal rearrangement errors. NucMerge was compared to two existing alternatives, namely Metassembler and GAM-NGS.

**Conclusions:** The benchmarking results show that NucMerge has generally better performance than the other tools tested, providing accuracy improvement of more assemblies. NucMerge is freely available at https://github.com/uio-bmi/NucMerge under the MPL license.

## 1. Background

In spite of the major breakthroughs in sequencing technologies over the last ten years, genome assembly using second-generation sequencing reads still remains a complicated problem. This is mainly due to the repeated structure of genomes and specific characteristics of the second-generation sequencing technologies, such as a large volume of data, relatively short length of sequenced fragments, uneven sequencing coverage, and presence of sequencing errors and chimeric reads [1,2]. A great number of assemblers with different heuristic approaches have been developed to deal with these challenges. However, the obtained assemblies may still be fragmented and containing errors.

A widely used approach to get an assembly using second-generation sequencing reads is to run multiple assemblers with different parameters and choose the best solution, for example based on assembly statistics, or agreement with physical, genetic, or optical maps. The rationality of this strategy has been confirmed by several evaluations like GAGE and Assemblathon [3-6]. The studies have revealed that the performance of assemblers varies a lot across different organisms, sequencing data, and assembler parameter settings and that it is impossible to predict in advance which assembler should be used. However, the studies have also shown that no single assembler has outperformed other assemblers in all evaluation metrics used, meaning that the process of choosing the best assembly is always a trade-off between different criteria (e.g. number of errors versus contiguity, number of insertions and deletions versus number of structural rearrangements, and so on).

To avoid the dilemma of determining which assembly is the best, an alternative approach would be to combine several assemblies into one. This process, called assembly reconciliation or assembly merging, enables the exploitation of advantages of different assemblers, resulting in a more contiguous solution with fewer errors. Existing merging tools can be divided into three groups: tools developed to correct errors and close internal gaps (e.g. Reconciliator [7]), tools aiming at the extension of assembly fragments and closure of gaps between fragments (e.g. Mix [8], GARM [9], ZORRO [10], and CISA [11]), and tools intended to improve both the accuracy and contiguity of a final assembly (e.g. GAA [12], GAM-NGS [13], and Metassembler [14]).

The tools aimed at assembly accuracy improvement (1) detect conflicting regions between alternative assemblies, (2) exploit different analysis, such as the compression-expansion statistic (CE statistic) [7] or depth-of-coverage analysis, to detect inconsistencies in the alignments of reads mapped back to the assemblies, and (3) choose the most reliable alternatives. The analysis based on the CE statistic has been first implemented in the Reconciliator tool and then incorporated in GAA and Metassembler. The CE statistic compares the mean insert size of the reads that span a particular region against the mean insert size of all reads. The depth-of coverage analysis has been used in GAM-NGS. It compares the read coverage of a particular region against the global read coverage. Although there has been some development in the field, there still seems to be possibilities for improvement.

In this article, we present a new assembly merging tool called NucMerge. It uses NucDiff [15] to detect differences between two alternative assemblies, and it uses Pilon [16] and NucBreak [17] to evaluate the correctness of assembly sequences in its pipeline. Pilon detects structural and local assembly errors of different lengths by analysing discordant paired-end read alignments and abnormal coverages. NucBreak detects structural errors and local errors longer than 30 bp in assemblies. In contrast to Pilon, it does this by analysing the alignments of reads that are properly mapped to an assembly (where both reads in a pair are fully aligned in correct orientation at a reasonable distance) and exploiting information about alternative alignments of the reads [17]. Being based on different methods, the tools complement each other, enabling the correction of more errors than either tool corrects alone.

NucMerge enables correction of inversions and local errors (insertions, deletions, and substitutions) and localization of rearrangement errors in genome assemblies. The tool has been compared with GAM-NGS and Metassembler. Only the ability of the tools to improve accuracy of assemblies has been studied. The results have shown that the error detection approach used in NucMerge is more effective than the CE-statistics and depth-of-coverage analysis, allowing NucMerge to improve assemblies in more cases compared to the other tools.

### 2. Implementation

NucMerge has been developed to improve the quality of an assembly, called the target assembly, by the correction of local errors and inversions and by the splitting of assembly fragments in the regions where inter- and intra-chromosomal rearrangements are located. Error correction and sequence splitting are performed based on information provided by another available assembly, called the query assembly, and a set of Illumina paired-end reads from the same genome.

### 2.1 Method description

The NucMerge pipeline consists of the evaluation, comparison and correction steps (see Figure 1). The evaluation and comparison steps prepare information needed for the correction step by running several external tools. During the evaluation step, NucBreak and Pilon analyse the target and query assemblies separately using information in the paired-end reads and predict erroneous regions of the assemblies. At the comparison step, NucDiff compares the query and target assemblies and outputs the types and locations of all differences between them.

**Figure 1.**
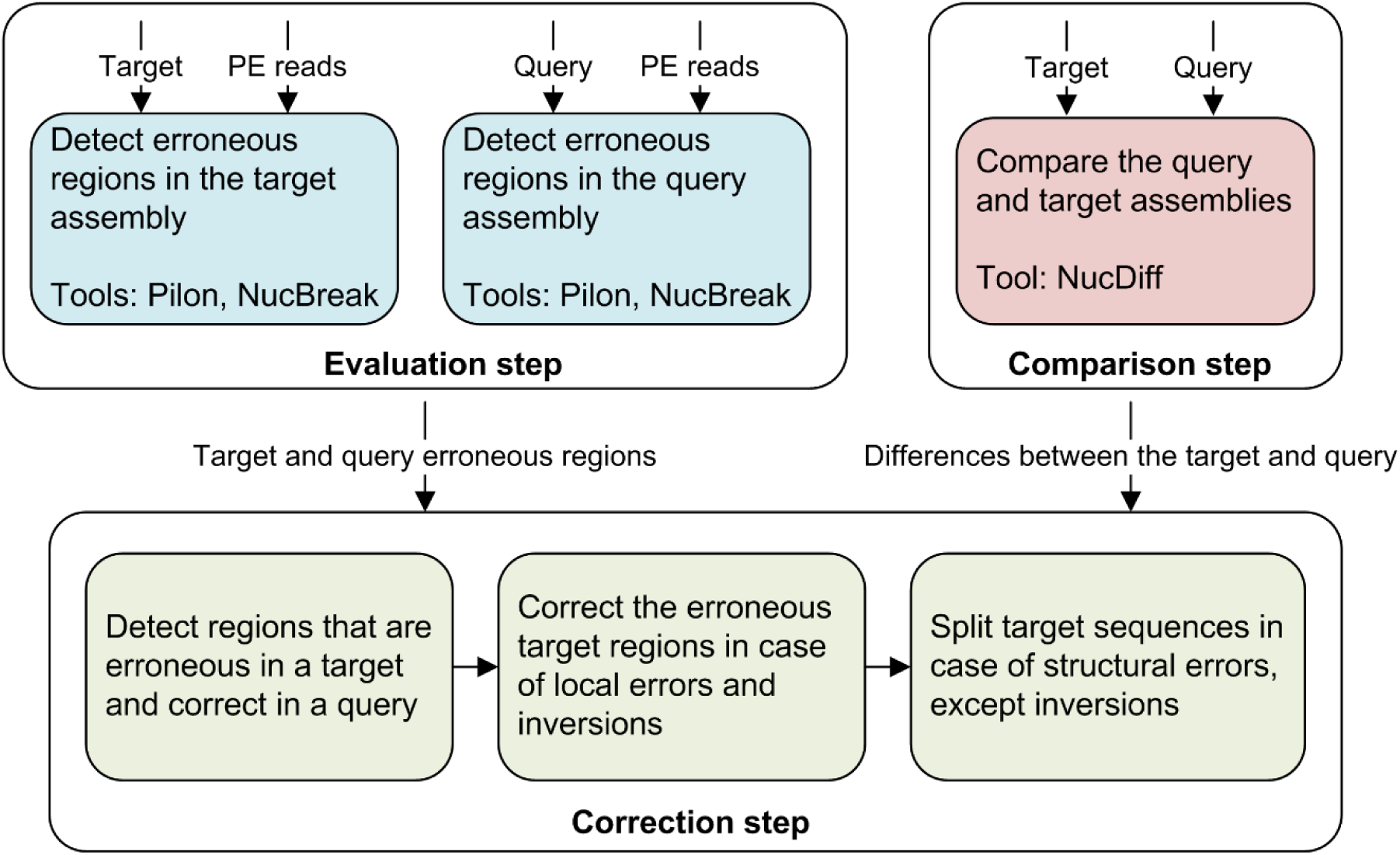
NucMerge workflow

Having collected all necessary information from the first two steps, NucMerge continues with the correction step. First, it detects erroneous target assembly regions suitable for alteration. For this purpose, it identifies differences located in the regions that have been predicted to contain errors in the target assembly by NucMerge and/or Pilon and have been predicted by the same tool(s) to be error free in the query assembly. Then NucMerge corrects or splits target assembly sequence regions containing the differences found, depending on the types of differences. If a target assembly region is related to an insertion, deletion, substitution or inversion difference, the region is substituted with the corresponding query assembly region. If a target assembly region is related to any inter- and intra-chromosomal rearrangement difference, the target assembly sequence is split in this region. The final result is an assembly with improved accuracy and a file with information about all changes performed.

### 2.2 Datasets

For testing purposes, we created three different datasets. The first dataset was created on the base of the data provided by the Assemblathon 1 project [5]. An artificially evolved human chromosome 13 (hg18/NCBI36), simulated Illumina paired-end read library with 40x coverage, and three genome assemblies obtained by Velvet [18], Meraculous [19], and AllPaths-LG [20] (named as C2, G1, and Q1, respectively, in the Assemblathon 1 project and in this article) were downloaded from the Assemblathon 1 website [21].

The second dataset consisted of 8 bacterial genomes, MiSeq Illumina paired-end read libraries provided for these genomes, and assemblies generated using the ABySS (version 2.0.2) [22], SPAdes (version 3.11.0) [23] and Velvet (version 1.2.10) assemblers. These assemblers were chosen because they are popular and were readily available. The genomes of the *Bordetella pertussis* str. J081, *Brucella melitensis* str. 1, *Enterobacter cloacae* str. AR_0136, *Escherichia coli* str. 2014C-3599, *Klebsiella pneumoniae* str. SGH10, *Pseudomonas aeruginosa* str. AR_0095, *Salmonella enterica* str. CFSAN047866, and *Staphylococcus aureus* str. CFSAN007896 organisms were downloaded from the NCBI database [24]. The reads were downloaded from the EBI database [25]. The genomes accession numbers and information about the read libraries are given in [Additional file 1: Table S1]. The parameter settings used to run ABySS, SPAdes and Velvet are described in [Additional file 1].

The third dataset was created on the basis of the data provided by the GAGE B project [4]. We used assemblies generated by the ABySS, CABOG [26], MaSuRCA [27], SGA [28], SOAPdenovo [29], SPAdes, and Velvet assemblers based on MiSeq Illumina reads from *Bacillus cereus* ATCC 10987 and HiSeq Illumina reads from *Rhodobacter sphaeroides* 2.4.1 and *Vibrio cholerae* CO 1032(5). The assemblies and reads were downloaded from the GAGE-B official website [30]. For evaluation, we used *B. cereus* ATCC 10987 (accession GCA_000008005.1), *R. sphaeroides* 2.4.1 (accession GCA_000012905.2), and *V. cholerae* O1 biovar eltor str. N16961 (accession GCA_000006745.1) as reference genomes. They were downloaded from the NCBI database.

## 3. Results

In this section, we explore the ability of NucBreak, Metassembler, and GAM-NGS to produce a new assembly with fewer errors using target and query assemblies and Illumina paired-end reads.

We excluded the GAA and Reconciliator tools from the testing. GAA had already been compared to GAM-NGS in [13] and [31]. In both cases, GAA showed its predisposition to introduction of structural rearrangements and duplication errors. Reconciliator is no longer maintained and not available any more. We also excluded MIX, Zorro, and CISA, since they only elongated the assembly sequences and filled in gaps but did not correct local assembly errors. In all tests performed, the tools were run with their default settings, except for the minimum and maximum insert size parameters, which should be specified for Metassembler and GAM-NGS. The exact parameter values used in each test are given in [Additional file 1].

To evaluate the results, we compared the initial and resulting assemblies to the reference genomes using the dnadiff utility [32] and assessed the accuracy of the assemblies based on the three following metrics: (1) the total number of bases involved in insertions, deletions and substitution errors (noted as the length of local errors), (2) the total number of inter- and intra-chromosomal translocations and inversions (noted as the number of structural errors), and (3) the total number of alignment breakpoints occurring due to the presence of long insertion and deletion errors as well as structural errors (noted as the number of breakpoints).

In order to improve visualization of the results, we show the differences between the values of the new assemblies and initial assemblies instead of their absolute values. In some figures, a logarithmic scale is applied.

### 3.1 Pairwise assembly merging validation using simulated datasets

First, we explored the ability of NucBreak, Metassembler, and GAM-NGS to correct errors when simulated data was given. For this purpose, we used an artificially evolved diploid genome, simulated reads, and three assemblies C2, G1, and Q1 generated by the Velvet, Meraculous, and AllPaths-LG assemblers respectively (see Section 2.2, the first dataset for details). We ran the tools for the following pairs of assemblies: C2 and G1, C2 and Q1, G1 and C2, and G1 and Q1, where the first assembly in a pair was a target assembly and the second assembly was a query assembly. The results for the two other combinations of assemblers were not included because their analysis with dnadiff was not completed within two weeks. The newly obtained assemblies were then compared with the reference genome using dnadiff. The comparison results are given in Figures 2-4.

**Figure 2.**
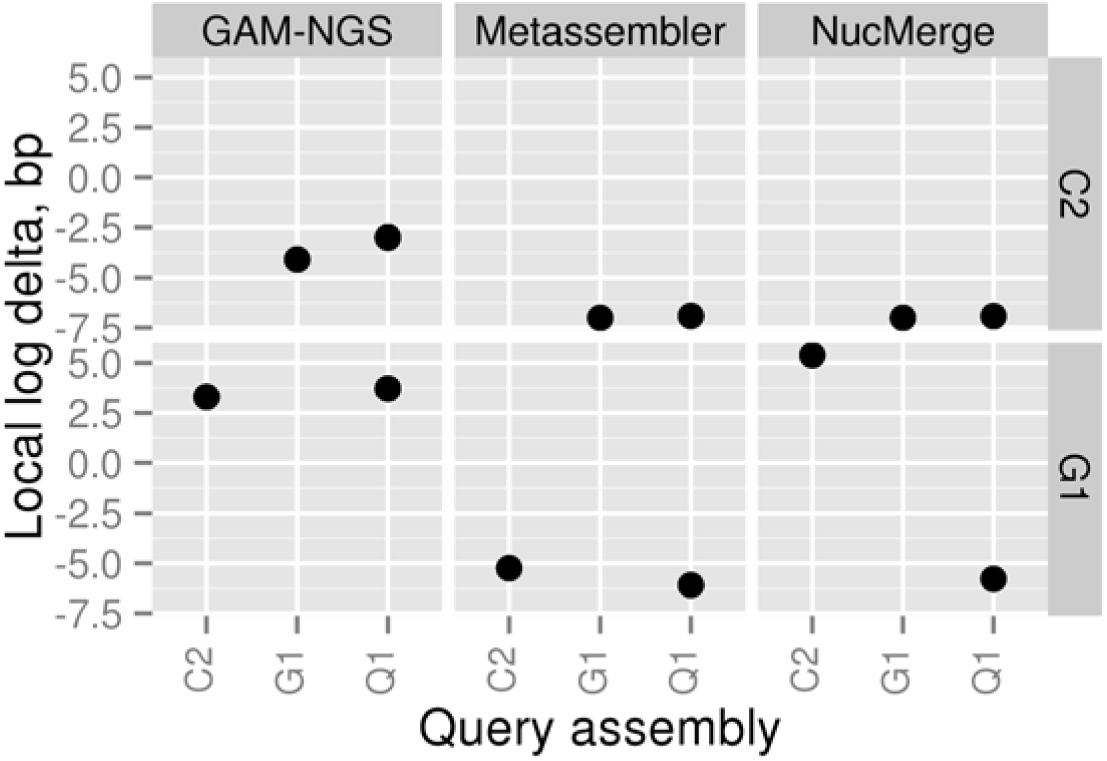
Differences between the total lengths of local errors in the initial (target) and final assemblies when the simulated dataset is used. All values are given in a logarithmic scale. A negative number means that the given tool has decreased the number of bases involved in local errors in the final assembly compared to the initial one. The C2, G1, and Q1 assemblies correspond to assemblies obtained using Velvet, Meraculous, and AllPaths-LG, respectively.

**Figure 3.**
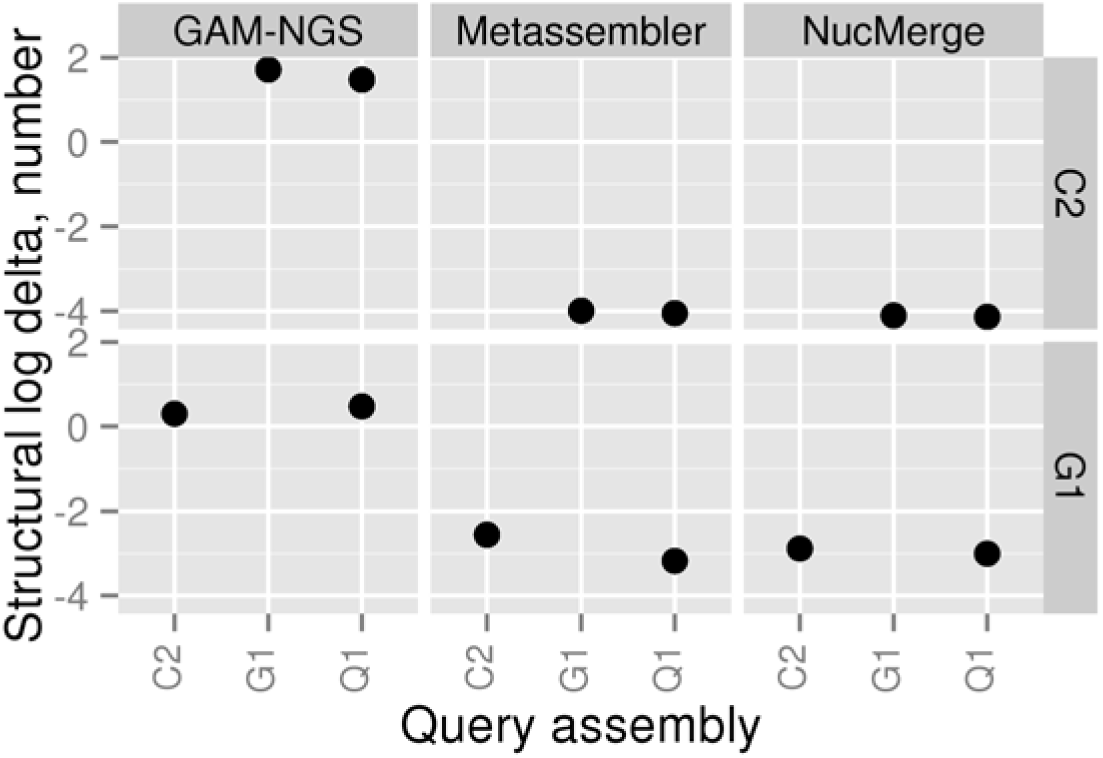
Differences between the number of structural errors in the initial (target) and final assemblies when the simulated dataset is used. All values are given in a logarithmic scale. A negative number means that the given tool has decreased the number of structural errors in the final assembly compared to the initial one. The C2, G1, and Q1 assemblies correspond to assemblies obtained using Velvet, Meraculous, and AllPaths-LG, respectively.

**Figure 4.**
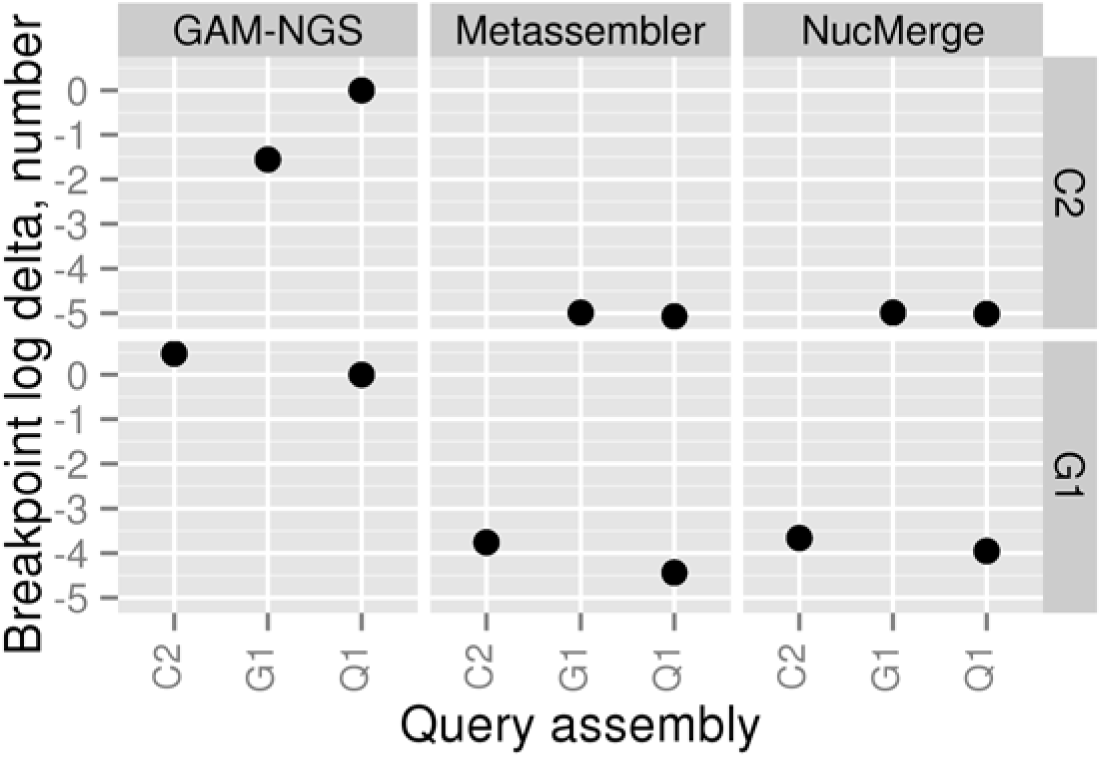
Differences between the number of breakpoints in the initial (target) and final assemblies when the simulated dataset is used. All values are given in a logarithmic scale. A negative number means that the given tool has decreased the number of breakpoints in the final assembly compared to the initial one. The C2, G1, and Q1 assemblies correspond to assemblies obtained using Velvet, Meraculous, and AllPaths-LG, respectively.

As seen from Figures 2-4, all tools managed to generate improved assemblies according to the different metrics studied. Metassembler obtained new assemblies with a decreased length of local errors and a reduced number of structural errors and breakpoints in all four cases. Best results were obtained with G1 as target assembly and Q1 as query assembly. In this case, the length of local errors was lowered by more than one million bases, and the number of structural errors and breakpoints was reduced by 1502 and 27440, respectively.

NucMerge performed very similar to Metassembler in all cases, except for the one case with G1 as target assembly and C2 as query assembly. In this case, NucMerge increased the length of local errors by 244436 bases while reducing the number of structural errors and breakpoints similarly to Metassember.

In contrast to the other tools, GAM-NGS performance varied considerably depending on the cases tested and metrics studied. It enabled the reduction of the length of local errors in two cases and the number of breakpoints in one case. It decreased the number of structural errors in none of the cases. None of the assemblies generated by GAM-NGS had all values reduced for the evaluated metrics.

### 3.2 Pairwise assembly merging validation using real datasets

Next, we explored the ability of NucBreak, Metassembler, and GAM-NGS to correct errors when real data was given. We downloaded reads for eight bacterial genomes and generated assemblies by using ABySS, SPAdes, and Velvet (see Section 2.2, the second dataset for full description of the data and assembler parameter settings used). NucBreak, Metassembler, and GAM-NGS were run for all possible pairs of assemblies separately for each organism. The obtained assemblies were then compared with reference genomes using dnadiff. The comparison results are presented in Figures 5-7.

**Figure 5.**
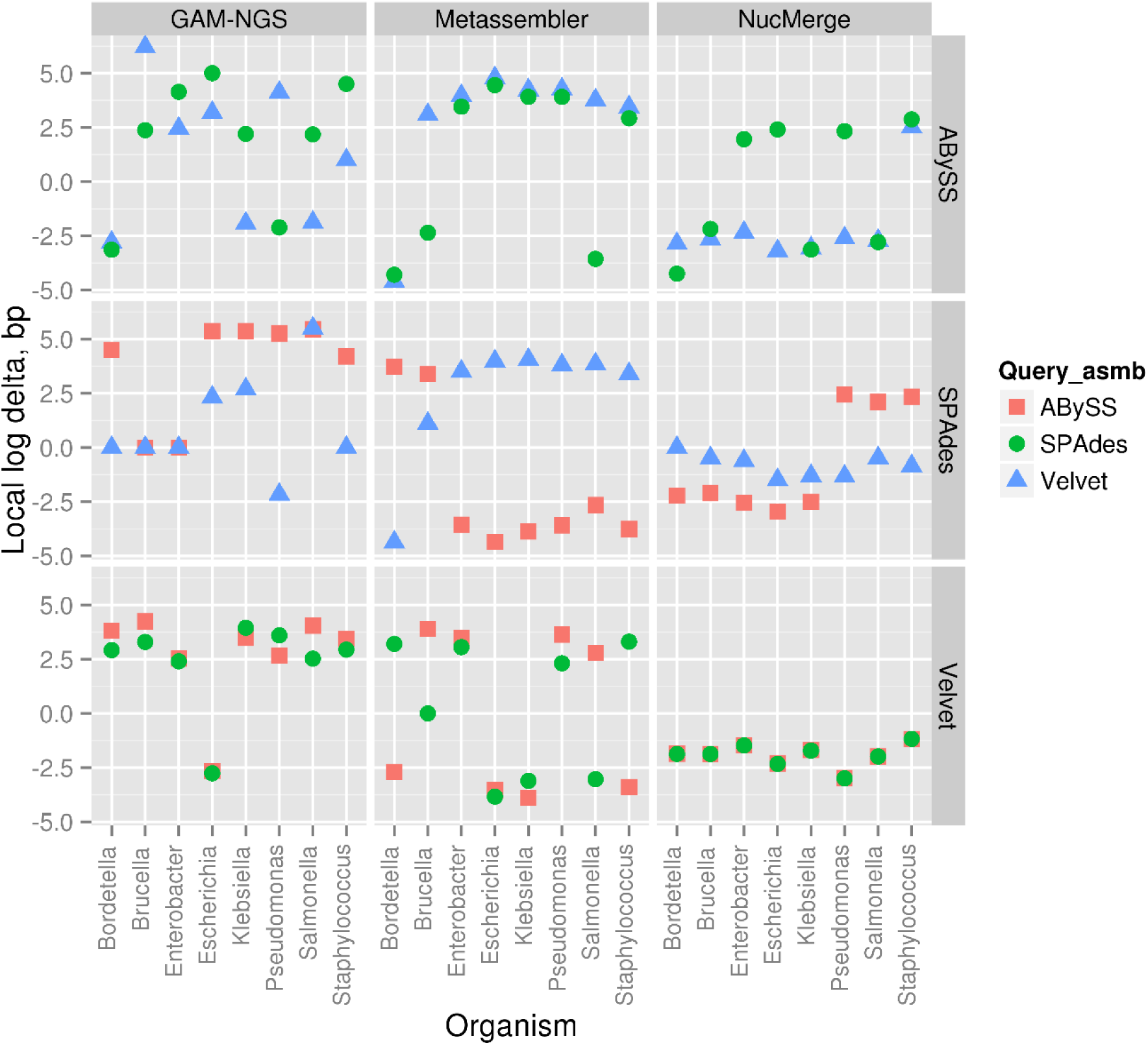
Differences between the total lengths of local errors in the initial (target) and final assemblies when the indicated bacterial genomes in the real dataset are used. All values are given in a logarithmic scale. A negative number means that the given tool (columns) has decreased the number of bases involved in local errors in the final assembly compared to the initial one. Query assemblies (Query_asmb) are indicated with colours. The row names denote the target assemblies.

**Figure 6.**
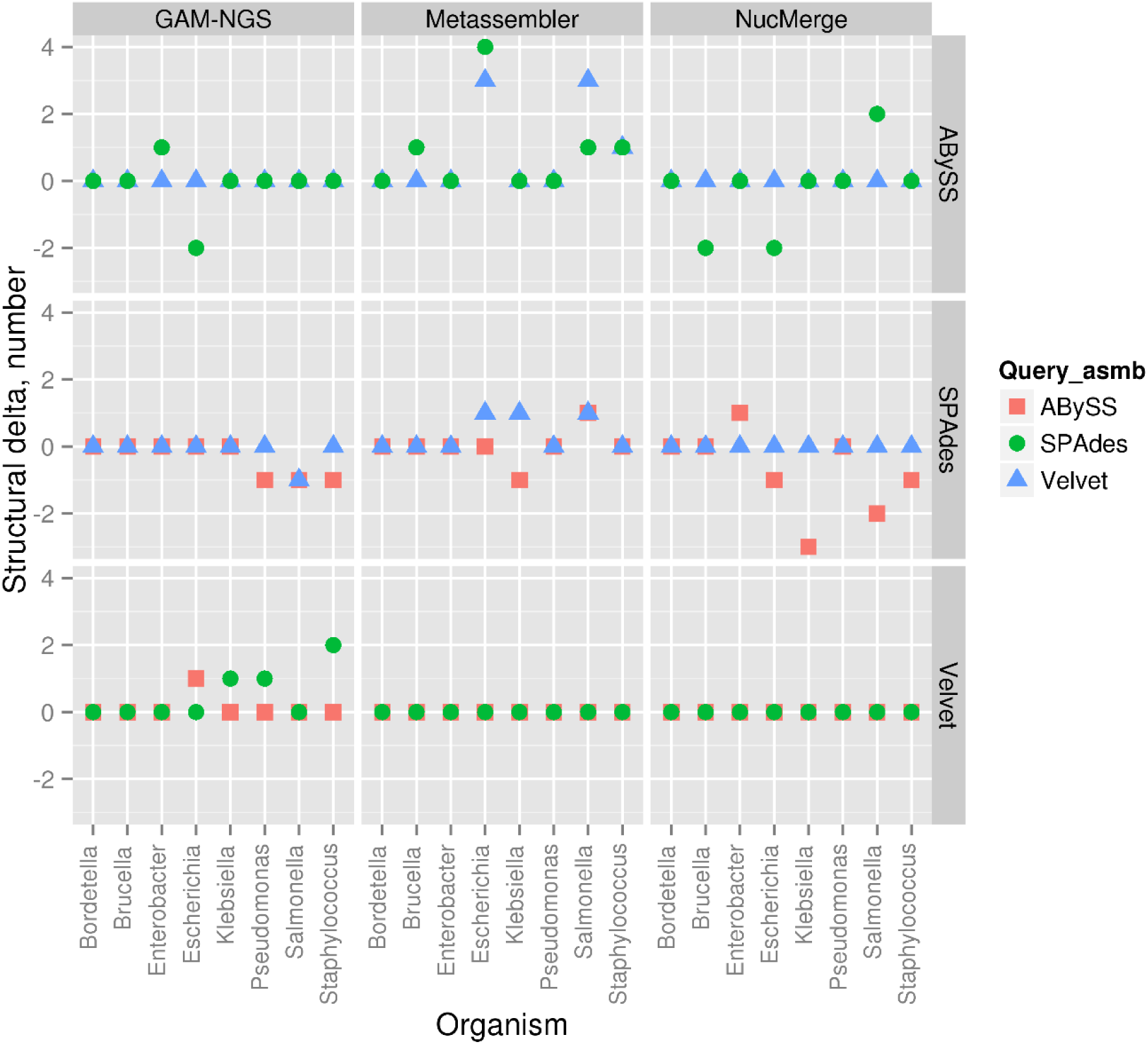
Differences between the number of structural errors in the initial (target) and final assemblies when the indicated bacterial genomes in the real dataset are used. A negative number means that the given tool (columns) has decreased the number of structural errors in the final assembly compared to the initial one. Query assemblies (Query_asmb) are indicated with colours. The row names denote the target assemblies.

**Figure 7.**
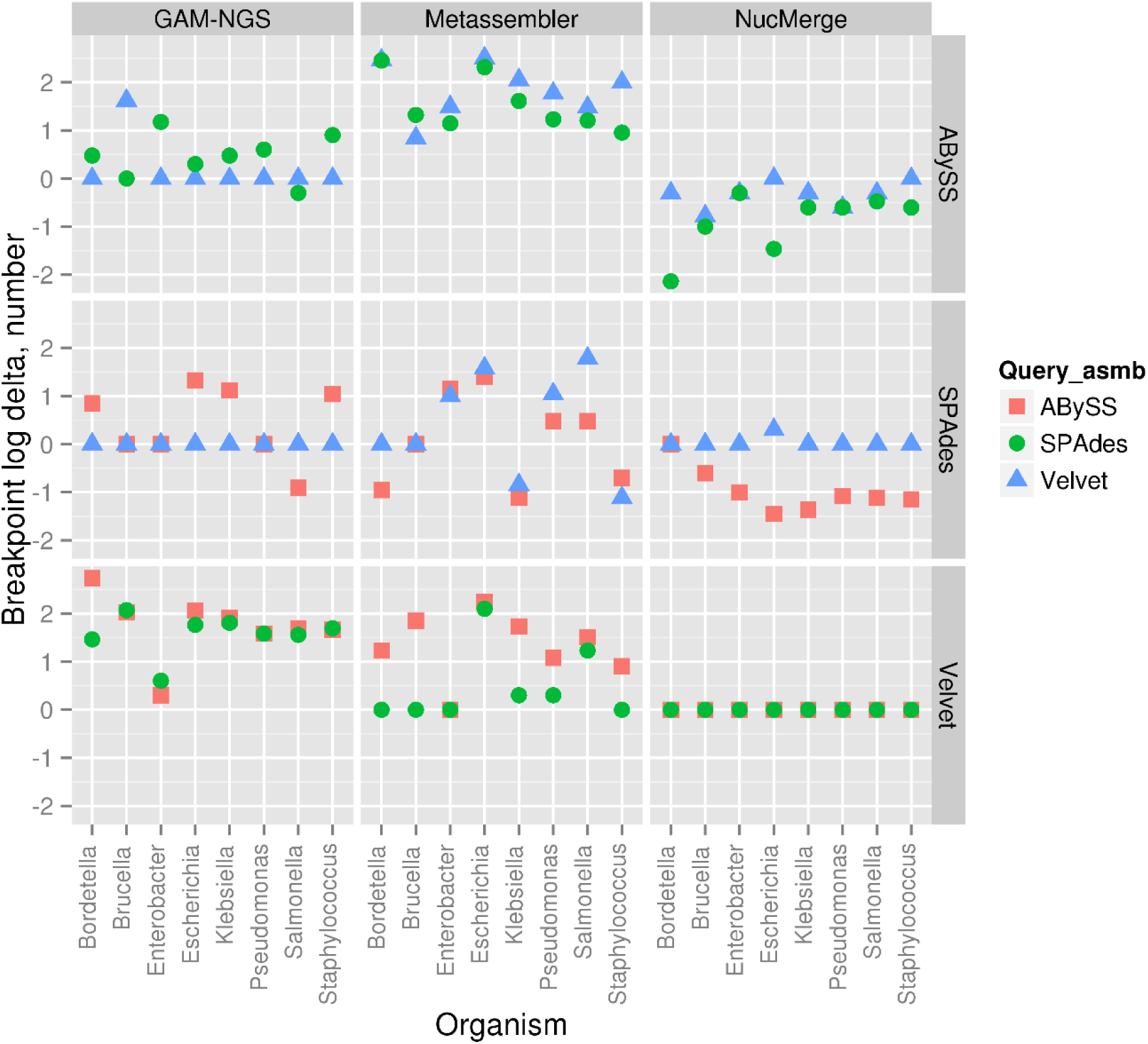
Differences between the number of breakpoints in the initial (target) and final when the indicated bacterial genomes in the real dataset are used. All values are given in a logarithmic scale. A negative number means that the given tool (columns) has decreased the number of breakpoints in the final assembly compared to the initial one. Query assemblies (Query_asmb) are indicated with colours. The row names denote the target assemblies.

As shown in Figures 5-7, the performance of all tools varied a lot depending on the data used and the metrics studied. However, all three tools were able to generate new assemblies where all metrics were decreased or remained the same compared to the initial target assembly. Such improved assemblies were observed in 36 cases for NucMerge, in 3 cases for Metassembler and in 4 cases for GAM-NGS out of 48 cases. The assemblies, where one or more metrics were increased, were observed in 12 cases for NucMerge, 44 cases for Metassembler and 38 cases for GAM-NGS. In the remaining cases, the metrics were unchanged.

NucMerge demonstrated its better performance by decreasing values for each metric in many more cases than the other tools. Metassembler reduced the length of local errors in more cases than GAM-NGS, while GAM-NGS reduced the number of structural errors and breakpoints in more cases than Metassembler.

### 3.3 Multiple assembly merging validation

Finally, we analysed the accuracy of the NucMerge, GAM-NGS and Metassembler solutions generated by the sequential merging of a target assembly with several query assemblies. To perform this, we downloaded reads and assemblies generated by the ABySS, CABOG, MaSuRCA, SGA, SOAPdenovo, SPAdes, and Velvet assemblers for three bacterial genomes (see Section 2.2, the third dataset for full description of the data used). We started the merging process with the assemblies with the longest and the second longest N50 (the SOAPdevo and CABOG assemblies for *B. cereus*, the MaSuRCA and SPAdes assembly for *R. sphaeroides* and the MaSuRCA and ABySS for *V. cholerae*) as target assemblies and applied the other assemblies as query assemblies in three different orders: (1) in order of decreasing N50 values, (2) in order of increasing N50 values and (3) in a random order. The exact merging order for each test is given in Table 1. The final solutions were then compared with reference genomes using dnadiff. The comparison results are presented in Figures 8-10.

**Table 1:** Merging order of the assemblies for each test performed. So, Ca, Ms, Ve, Sp, Ab, and Sg denotes the SOAPdenovo, CABOG, MaSuRCA, Velvet, SPAdes, ABySS, and SGA assemblies, respectively. The first assembly in the order sequence is a target assembly, all others are query assemblies. In the table cell, the first order sequence corresponds to the one where the target assembly has the longest N50, while the second order sequence corresponds to the one where the target assembly has the second longest N50.

| Reference | N50 decreasing | N50 increasing | Random, test 1 | Random, test 2 | Random, test 3 |
| --- | --- | --- | --- | --- | --- |
| *B. cereus* | SoCaMsVeSpAbSg | SoSgAbSpVeMsCa | SoAbMsSpVeSgCa | SoSpCaVeAbMsSg | SoCaSpAbSgVeMs |
|  | CaSoMsVeSpAbSg | CaSgAbSpVeMsSo | CaSpSgSoMsVeAb | CaAbSoSgVeMsSp | CaSpMsAbVeSgSo |
| *R. sphaeroides* | MsSpVeCaSoAbSg | MsSgAbSoCaVeSp | MsCaSpSoAbVeSg | MsVeSoSgAbCaSp | MsSoSpAbVeSgCa |
|  | SpMsVeCaSoAbSg | SpSgAbSoCaVeMs | SpSgSoVeAbCaMs | SpMsSgAbVeCaSo | SpVeSgMsSoCaAb |
| *V. cholerae* | MsAbSoVeSpCaSg | MsSgCaSpVeSoAb | MsAbSgSpSoVeCa | MsSoCaSgAbSpVe | MsVeSoCaSpAbSg |
|  | AbMsSoVeSpCaSg | AbSgCaSpVeSoMs | AbSoVeSgCaMsSp | AbCaMsSoSgSpVe | AbVeCaSoMsSgSp |

**Figure 8.**
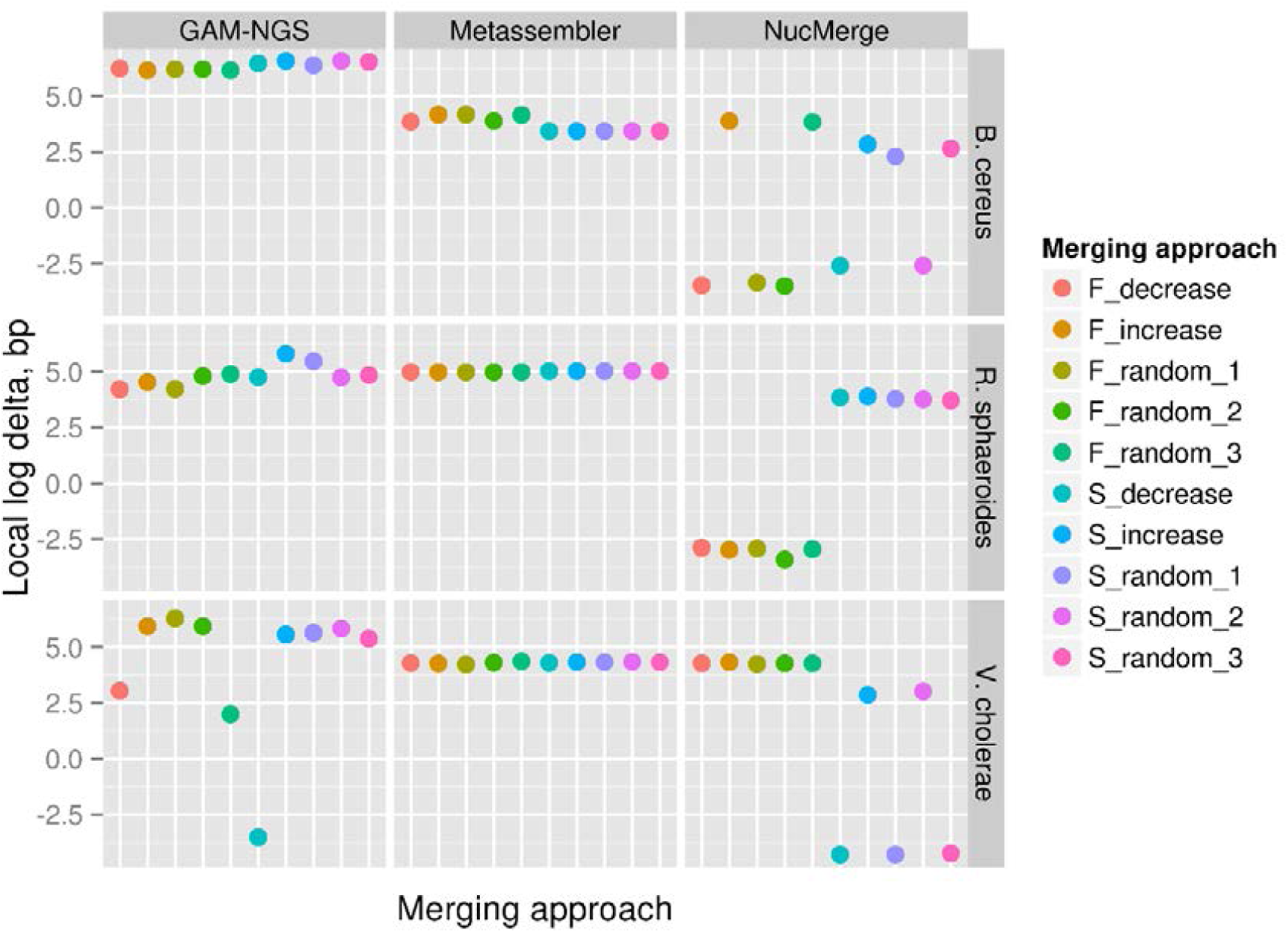
Differences between the total lengths of local errors in the initial (target) and final assemblies when the multiple assembly merging approach is applied. All values are given in a logarithmic scale. A negative number means that the given tool has decreased the number of bases involved in local errors in the final assembly compared to the initial one. In the notations of the merging approach names, F and S denote the cases where the target assembly with the longest N50 and second longest N50 are used, respectively. The labels “decrease”, “increase”, “random_1”, “random_2”, and “random_3” denote merging strategies and correspond to the N50 decreasing, N50 increasing and random test1, test2 and test3 merging strategies described in Table 1.

**Figure 9.**
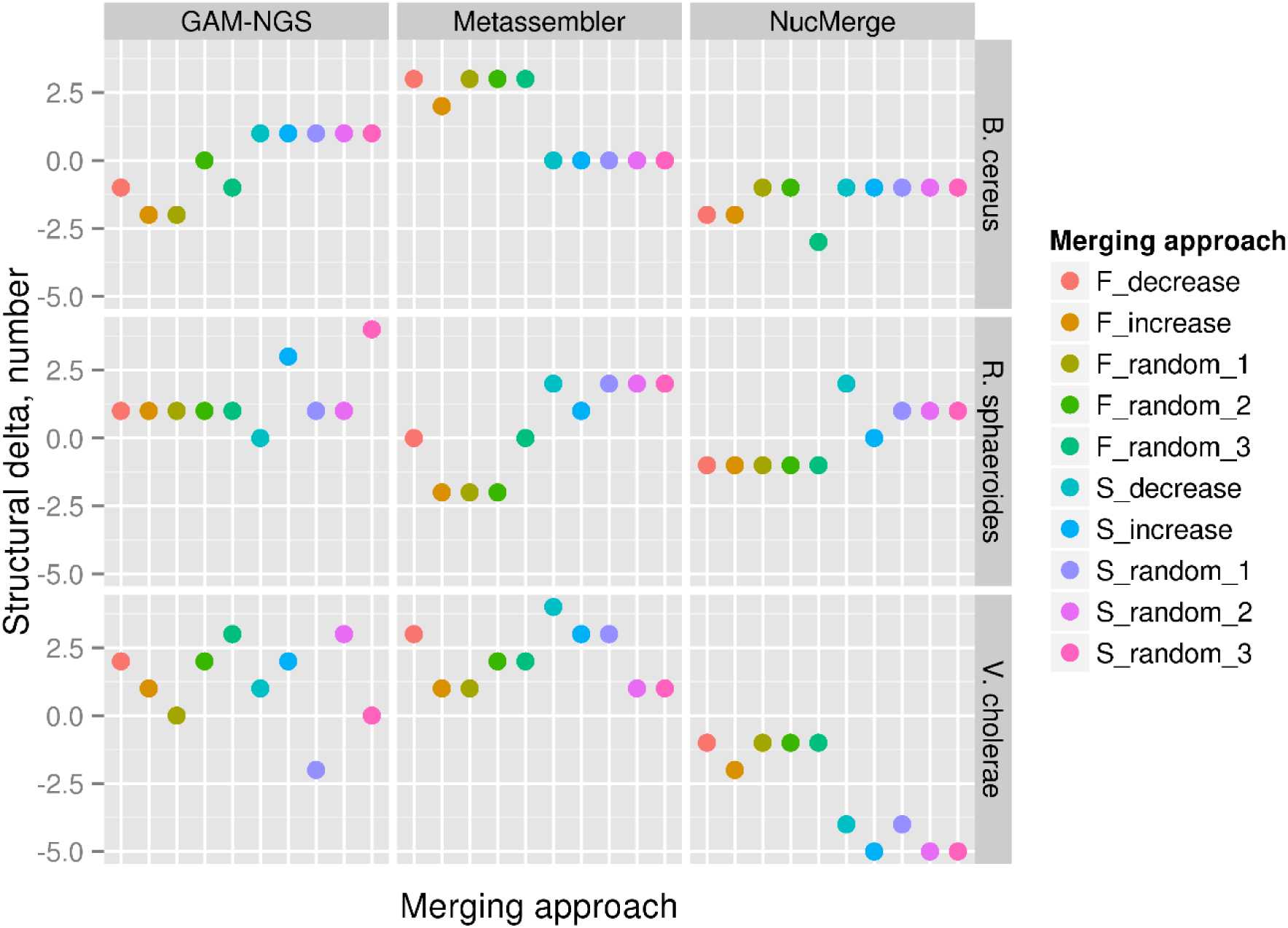
Differences between the number of structural errors in the initial (target) and final assemblies when the multiple assembly merging approach is applied. A negative number means that the given tool has decreased the number of structural errors in the final assembly compared to the initial one. In the notations of merging approach names, F and S denote the cases where the target assembly with the longest N50 and second longest N50 are used, respectively. The labels “decrease”, “increase”, “random_1”, “random_2”, and “random_3” denote merging strategies and correspond to the N50 decreasing, N50 increasing and random test1, test2 and test3 merging strategies described in Table 1.

**Figure 10.**
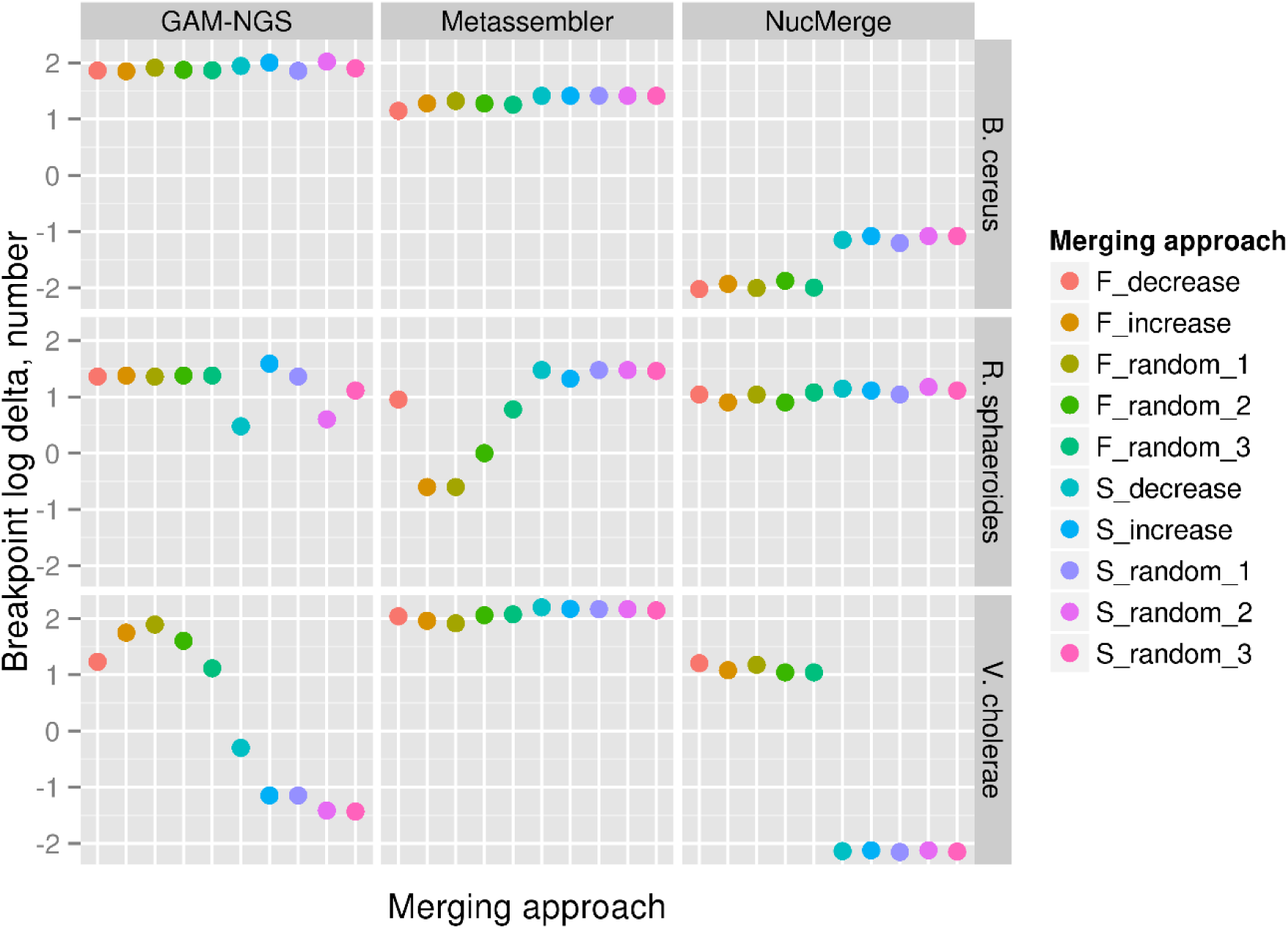
Differences between the number of breakpoints in the initial (target) and final assemblies when the multiple assembly merging approach is applied. All values are given in a logarithmic scale. A negative number means that the given tool has decreased the number of breakpoints in the final assembly compared to the initial one. In the notations of merging approach names, F and S denote the cases where the target assembly with the longest N50 and second longest N50 are used, respectively. The labels “decrease”, “increase”, “random_1”, “random_2”, and “random_3” denote merging strategies and correspond to the N50 decreasing, N50 increasing and random test1, test2 and test3 merging strategies described in Table 1.

From the plots presented in Figures 8-10, it can be concluded that NucMerge has overall better performance than the other tools according to the metrics evaluated. It decreased the length of local errors and number of breakpoints approximately in half of the cases and reduced the number of structural errors in the most cases. It also generated 8 out of 30 new assemblies where all metrics were reduced. In contrast, Metassembler and GAM-NGS increased all metrics in almost all cases. They did not produce any new assembly where all metrics were improved.

The comparison of the results obtained with different merging strategies showed that the strategy where assemblies were added in order of decreasing N50 values was beneficial for local error correction. The strategy where assemblies were added in order of increasing N50 values showed good results when it was applied for structural error detection. The merging strategy with random assembly order showed both good and bad results for all metrics and all tools. The effectiveness of each strategy to decrease the number of breakpoints varied depending on tools, organisms, and assembly pairs.

## 4. Discussion

In this work, we have introduced a tool called NucMerge. With the assistance of alternative assemblies, NucMerge corrects inversions and local errors (such as insertions, deletions, and substitutions) and locates structural inconsistencies (such as inter- and intra-chromosomal rearrangements) in assemblies. It is based on the usage of two existing tools for assembly error detection, called Pilon and NucBreak. These tools are utilized as an alternative to the CE-statistics and depth-of-coverage analysis exploited in the existing tools, such as GAM-NGS and Metassembler. The results show that the combination of methods, as it is implemented in NucMerge, in general is more beneficial for assembly merging than a single method, enabling production of improved assemblies. For bacterial genomes, NucMerge performed better in most cases, while for the artificial genome it performed equal to the best of the tools, except for one case.

The proposed approach does not completely eliminate the production of assemblies with deteriorated accuracy. According to the results, NucMerge has also generated new assemblies with more structural errors and longer length of local errors in some cases. To enable manual validation of all modifications proposed and reduce the number of false modifications, NucMerge outputs information about the locations of structural inconsistencies detected and modifications performed in addition to the modified assembly sequences. The information is stored in GFF3 format [33] and can be easily uploaded to a genome browser, e.g. IGV [34,35], together with the assembly and read alignments.

During the testing, we explored both pairwise and multiple assembly merging approaches. The results obtained for the pairwise merging did not reveal any dependencies between the successful merging and certain combinations of assemblers for all tools tested. The same pair of assemblers produced the assemblies with both improved and deteriorated accuracy for different organisms.

During the multiple merging testing, we merged sequentially target assemblies with other assemblies in five different orders. The orders included three different random combinations and two combinations depending on the N50 values (one in increasing N50 order and one in decreasing N50 order). Although the random query assembly order gave rise to the solutions with the best accuracy compared to the other merging strategies, it was also associated with the cases of the strongest accuracy degradation. Not having any possibilities to assess the quality of the obtained assemblies, it is clearly a risky strategy. The strategy with the decreasing N50 query assembly order always produced solutions with one of the lowest number of bases involved in local errors compared to the other merging strategies for all tools. The same strategy was beneficial for reduction of structural errors for GAM-NGS. However, for NucMerge and Metassembler, the best strategy to reduce the structural error number was the one with the increasing N50 query assembly order.

The current version of the tool described here focuses on the matter of assembly accuracy improvement only. However, we also intend to extend NucMerge’s functionality by merging assembly sequences and resolving rearrangements errors to improve assembly contiguity.

## 5. Conclusion

We have presented NucMerge, a new tool for merging two or more alternative assemblies into a new assembly with improved accuracy with the assistance of paired-end Illumina reads. The tool enables correction of inversions and local errors, such as insertions, deletions and substitutions, and provide locations of structural errors, such as inter- and intra-chromosomal rearrangements, in a target assembly. In contrast to the other existing tools, NucMerge exploits several methods for assembly error detection, such as analysis of discordant paired-end read alignments, abnormal coverages, alternative read alignments and properly mapped read pairs, by using Pilon and NucBreak in its pipeline. The tool was compared to the existing alternatives, namely Metassembler and GAM-NGS. The results have shown that it improves assemblies in more cases compared to the other tools using both pairwise and multiple assembly merging approaches.

## Supporting information

## Declarations

## Acknowledgements

The authors wish to thank the Centre for Ecological and Evolutionary Synthesis (CEES) for access to the computational infrastructure (‘cod’ servers) that enabled the bioinformatics analysis for this project.

## Availability of data and materials

- Project name: NucMerge
- Project home page: https://github.com/uio-bmi/NucMerge
- Operating system(s): Unix-like system such as Ubuntu Linux and MacOS X.
- Programming language: Python
- Other requirements: Python 2.7
- License: Mozilla Public License (MPL), version 2.0
- Any restrictions to use by non-academics: No
- Additional data: all data used is available as described in Section 2.2

### Authors’ contributions

KK designed and implemented NucMerge. KK, GKS, AJN, and TR suggested the experiments performed. KK performed all the experiments. KK and TR wrote the manuscript. GKS and AJN revised the manuscript. All authors read and approved the final manuscript.

## Ethics approval and consent to participate

Not applicable.

## Consent for publication

Not applicable.

## Additional files

**Additional file 1** Supplementary materials (PDF 46KB)

Table S1 List of bacterial genomes

## Abbreviations

CE: compression-expansion
bp: base pairs

