## Supplementary material for "NucMerge: Genome assembly quality improvement assisted by alternative assemblies and paired-end Illumina reads"

#### *Supplementary materials*

Ksenia Khelik, Alexander Johan Nederbragt, Geir Kjetil Sandve, Torbjørn Rognes

#### Supplementary methods

##### 1. The Velvet, ABySS and SPAdes parameter settings used to obtain assemblies

SPAdes was run with the “-t 2 -k 33 --cov-cutoff 2” parameter settings.
ABySS was run with “k=64” parameter setting.
Velveth was run with k-mer length equal to 31.
Velvetg was run with “-ins_length 180 -scaffolding yes -min_contig_lgth 250 -cov_cutoff 5” parameter settings.

##### 2. The Metassembler and GAM-NGS insert size parameter setting

For the tests with Assemblathon 1 data sets, we have used the minimum insert size equal to 100 and the maximum insert size equal to 500 for all assembly pairs.

For the tests with eight bacterial genomes, we have used the minimum insert size equal to 35 and the maximum insert size equal to 1120 for all assembly pairs.

For the tests with GAGE B data sets, we have used the minimum insert size equal to 450 and the maximum insert size equal to 700 for all assembly pairs.

### Supplementary tables

**Table S1** List of bacterial genomes.

| Genome | Genome length, Mb | Accession number | Reads length, bp (first, second) | Coverage | Read library accession number |
|---|---|---|---|---|---|
| *Bordetella pertussis* str. J081 | 4,11 | GCA_002859625.1 | 250<br>250 | 32x | SRR5829829 |
| *Brucella melitensis* str. 1 | 3,30 | GCA_900236405.1 | 243 ± 28.8<br>243 ± 28.7 | 40x | ERR2192800 |
| *Enterobacter cloacae* str. AR_0136 | 5,04 | GCA_002204775.1 | 233 ± 34.9<br>233 ± 34.8 | 23x | SRR4025988 |
| *Escherichia coli* str. 2014C-3599 | 5,48 | GCA_003018935.1 | 236 ± 39.0<br>236 ± 38.8 | 60x | SRR1609862 |
| *Klebsiella pneumonia* str. SGH10 | 5,72 | GCA_002813595.1 | 146 ± 15.8<br>146 ± 15.7 | 32x | SRR5082357 |
| *Pseudomonas aeruginosa* str. AR_0095 | 6.82 | GCA_002997005.1 | 229 ± 38.2<br>229 ± 36.9 | 60x | SRR3242025 |
| *Salmonella enterica* str. CFSAN047866 | 4,81 | GCA_003073535.1 | 244 ± 27.3<br>244 ± 27.3 | 37x | SRR3272258 |
| *Staphylococcus aureus* str. CFSAN007896 | 2,86 | GCA_003031425.1 | 236 ± 41.8<br>236 ± 41.7 | 28x | SRR5912676 |
